# ECloudGen: Leveraging Electron Clouds as a Latent Variable to Scale Up Structure-based Molecular Design

**DOI:** 10.1101/2024.06.03.597263

**Authors:** Odin Zhang, Jieyu Jin, Zhenxing Wu, Jintu Zhang, Po Yuan, Haitao Lin, Haiyang Zhong, Xujun Zhang, Chenqing Hua, Weibo Zhao, Zhengshuo Zhang, Kejun Ying, Yufei Huang, Huifeng Zhao, Yuntao Yu, Yu Kang, Peichen Pan, Jike Wang, Dong Guo, Shuangjia Zheng, Chang-Yu Hsieh, Tingjun Hou

## Abstract

Structure-based molecule generation represents a significant advancement in AI-aided drug design (AIDD). However, progress in this domain is constrained by the scarcity of structural data on protein-ligand complexes, a challenge we term the Paradox of Sparse Chemical Space Generation. To address this limitation, we propose a novel latent variable approach that bridges the data gap between ligand-only and protein-ligand complexes, enabling the target-aware generative models to explore a broader chemical space and enhancing the quality of molecular generation. Drawing inspiration from quantum molecular simulations, we introduce ECloudGen, a generative model that leverages electron clouds as meaningful latent variables—an innovative integration of physical principles into deep learning frameworks. ECloudGen incorporates modern techniques, including latent diffusion models, Llama architectures, and a newly proposed contrastive learning task, which organizes the chemical space into a structured and highly interpretable latent representation. Benchmark studies demonstrate that ECloudGen outperforms state-of-the-art methods by generating more potent binders with superior physiochemical properties and by covering a significantly broader chemical space. The incorporation of electron clouds as latent variables not only improves generative performance but also introduces model-level interpretability, as illustrated in a case study designing V2R inhibitors. Furthermore, ECloudGen’s structurally ordered modeling of chemical space enables the development of a model-agnostic optimizer, extending its utility to molecular optimization tasks. This capability has been validated through a single-objective oracle benchmark and a complex multi-objective optimization scenario involving the redesign of endogenous BRD4 ligands. In conclusion, ECloudGen effectively addresses the Paradox of Sparse Chemical Space Generation through its integration of theoretical insights, advanced generative techniques, and real-world validation. The newly proposed technique of leveraging physical entities (such as electron clouds) as latent variables within a deep learning framework may prove useful for computational biology fields beyond AIDD.

## Introduction

Developing new drugs is one of the most effective ways for humans to combat diseases^1^. Molecular generation is the crown jewel of Artificial Intelligence-Driven Drug Discovery (AIDD), aiming to design novel and effective drugs^2^. However, non-target specific generation of molecules, as done by many earlierly developped molecular graph generation methods^3,4^, cannot satisfy all the rigorous demands of drug design in the real-world scenarios. Therefore, recent research has shifted towards constrained molecular generation, especially focusing on structure-based molecular generation (SBMG)^5^, in which AI models try to propose molecules that fits into the given protein targets. This paradigm shift can be mathematically represented as modeling *p*(*G*|*p*) instead of *p*(*G*), where *G* and *p* denotes molecule and protein, respectively. Training molecular generation methods can be seen as conducting likelihood maximization on these probabilities, and owing to recent advances in generative models such as diffusion models^6^ and large language models^7^ (LLMs), researchers now possessn ever more potent binder tools to manage these complex probabilities. As a result, a plethora of models have emerged in the SBMG field. For example, DiffBP^8^, DiffSBDD^9^, and TargetDiff^10^ have adopted the one-shot generation protocol using the diffusion model, directly denoising 3D molecules within protein pockets. Pkt2Mol^11^, ResGen^5^, and SurfGen^12^ utilize the auto-regressive generation protocol with geometric/geodesic neural networks, sequentially adding atoms within protein pockets. Meanwhile, Lingo3DMol^13^ and PrefixMol^14^ leverage the transformer architecture to enhance the learning of *p*(*G*|*p*) in protein pockets.

Although these models have achieved many promising results in designing lead compounds, with some even validated through wet-lab experiments^15,16^, a long-standing yet often overlooked issue persists: data scarcity. This issue is not severe in the arbitrary molecular generation task, i.e., learning *p*(*G*), where the training dataset encompasses ∼10^10^ molecules^17^. However, in SBMG tasks of learning *p*(*G*|*p*), this point becomes critical. Specifically, the training pairs of learning *p*(*G*|*p*) derived from known binding complex structures, where are only ∼10^5^ unique molecules^18^ (**Figure 1A left**). This number is minuscule compared to the vast chemical libraries accessible (**Figure 1A middle**), not to mention the entire estimated chemical space (10^60^)^19^ of drug-like compounds (**Figure 1A right**). The **Figure 1A left** illustrates how sampling ‘holes’ in areas of low chemical space density can lead to poor quality in generated molecules. Herein, we formally characterize the data scarcity in the SBMG field as the ‘Paradox of Sparse Chemical Space Generation.’ This paradox highlights that although the primary goal of generative models is to explore and sample broader chemical spaces, the currently available protein-ligand pairs dataset confines them to a much narrower space, thus hampering the full potential of generative AI models.

**Figure 1.**
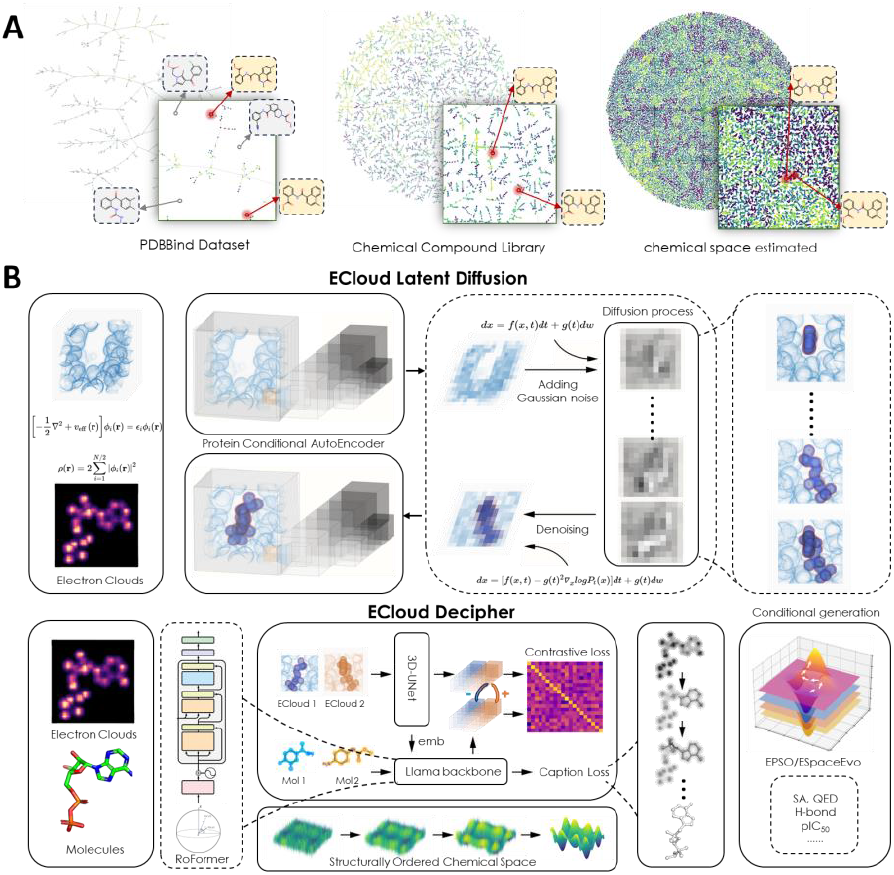
**A)** Illustration of the **Sparse Chemical Generation Space Paradox**, contrasting three scales of chemical spaces. The PDBBind dataset encompasses roughly 10^5^ protein-ligand pairs, the chemical compound library contains about 10^9^ molecules, and the vast estimated total chemical space holds approximately 10^60^ compounds. **B)** ECloudGen main modules. The first module generates learns *p*(*C*|*p*), which generates electron clouds from protein pockets; and the second module learns *p*(*G*|*C, p*) which translates these electron clouds into molecular ball-and-stick represenations.

As we all know, the quality and size of datasets are two essential ingredients for building succssful AI models. Therefore, it would be high rewarding if the proposed paradox can be reasonably overcome. To this end, we developed ECloudGen, which adopts a novel training technique combined with quantum physics to expand the chemical space in SBMG models, thereby fundamentally improving their generation quality. There are two major methodological considerations in ECloudGen. Mathematically, by identifying a latent variable that bridges protein-ligand pairs and ligand-only data, we can effectively integrate a broader chemical space from ligand-only data into the model. Specifically, we decompose the learning of *p*(*G*|*p*) to the learning of *p*(*G*|*C, p*) · *p*(*C*|*p*), where *C* is some prescribed latent variable. This approach allows the model to learn *p*(*C*|*p*) from ligand-only data while learning *p*(*G*|*C, p*) from protein-ligand pairs, thus accessing the joint chemical spaces encomposed by the two data sources. Generally, it is beneficial to consider a latent variable that is physically meaningful, rather than purely artificial, to introduce interpretability and controllability into the model. Bearing this idea in mind, we identified electron clouds - a fundamental description of molecules - as the latent variable, also called generative agent in the following context. Additionally, learning on electron clouds offer additional benefits. For example, since all interatomic interactions are fundamentally electromagnetic in nature and governed by electron clouds^20^, this representation offers a unified perspective on molecular tasks. More discussions can be found in **Part S1**.

To implement the above intuitions, ECloudGen is built with two major components: the first component, modeling *p*(*C*|*p*), is designed to generate appropriate electron clouds conditioned on a given protein pocket and is trained on the CrossDock dataset^21^; the second component, modeling *p*(*G*|*C, p*), is designed to decode the electron clouds, along with a given protein pocket, into a molecular graph and is further trained on large chemical libraries^17^. The model is illustrated in **Figure 1B**. In terms of detailed architectures, we implemented modern techniques to distill the chemical space from extensive training data. To be more specific, we introduced a 3D Latent Diffusion^22^ to efficiently sample high-fidelity electron clouds in the first module, and a Llama-like^7^ architecture to decipher molecules in the second module. We also proposed the CEMP (Contrastive Electron Clouds Pre-training) to facilitate the modeling of a structurally ordered chemical space for better generalization and molecular optimization. Subsequent experiments demonstrated that ECloudGen accesses a broader chemical space compared to ten baseline models, effectively alleviating the identified generation paradox. Thanks to this capability, ECloudGen has been proven to generate molecules that bind tightly and possess improved molecular properties, as evidenced in comprehensive benchmarks. In practical applications, we have further demonstrated another advantage of our physically meaningful latent variable, electron clouds: the provision of model-level interpretability and its utility in drug discovery. Furthermore, we leveraged the structurally ordered properties of the ECloudGen chemical space to construct a model-agnostic module for both single-condition and multi-condition molecular optimization. The effectiveness of this module was showcased in an oracle benchmark and a drug design scenario involving five constraint conditions. Combined with modern AI techniques and physical insights, ECloudGen offers a unique approach to molecular generation.

## Results and Discussion

### ECloudGen Generates Drug-like Molecules on the Benchmark

The potential of a molecule to become a viable drug is determined by its binding affinity to the target and its drug-like molecular properties. Therefore, molecular generation models are primarily evaluated on these two aspects. Previous structure-based molecular generation models have been effective in designing molecules with strong binding affinities. However, these models still face limitations in terms of drug-likeness properties. While some methods have succeeded in generating molecules with improved properties through fragment-wise approaches^23,24^, these strategies have not addressed the underlying issue of poor molecular properties, namely, the Sparse Chemical Space Paradox. In this study, we demonstrate how ECloudGen’s access to a larger chemical space can significantly enhance the quality of the generated molecules.

We comprehensively compared ECloudGen against ten baseline models, including Pocket2Mol^11^, ResGen^5^, GraphBP^25^, FLAG^23^, FragGen ^24^, DiffBP^8^, DiffSBDD^9^, TargetDiff ^10^, Lingo3dMol^13^, PrefixMol^14^. These models were trained and tested using the CrossDock dataset with a protein similarity data split^21^. To evaluate the quality of generated molecules, we used the Vina score^26^ to estimate their binding affinities with corresponding protein targets. Moreover, we used QED^27^, SA^28^, Lipinski Rule of Five^29^, and LogP^30^ to assess the drug-likeness of the molecules. It is important to note that the Vina score, which is derived from the summation of individual atom/fragment energy contributions, inherently favors larger molecules. To counteract this bias, we adopt Ligand Binding Efficiency (LBE)^31^, a metric commonly used in computer-aided drug discovery, which normalizes the Vina score based on the number of atoms, i.e., 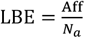, where Aff is the molecular binding affinity and *N*_*a*_ is the number of atoms. Previous studies have shown that LBE provides a more realistic measure than the unnormalized Vina score^32,33^. Hence, we adopted LBE as a better alternative for evaluating binding efficiency. More information of these metrics can be found in the **Method** section.

As drug design research typically focuses on a few molecules with the highest potential, we computed the average results for the top five molecules of each target in the test set. The results, summarized in **Table 1**, show that ECloudGen achieves the best results on the LBE metric, indicating its ability to design molecules that effectively bind to protein targets. Diffusion models, like DiffBP, DiffSBDD, and TargetDiff, exhibit relatively low performance on the QED metric. This may be due to these diffusion models’ adoption of a one-shot generation protocol, which attempts to estimate the likelihood *p*(*A*_*a*_, *E*_*a*_|*N*_*a*_), where *A*_*a*_, *E*_*a*_ are atomic type and bonding relationships of the molecules. This joint distribution has strong correlations; for instance, the type of an atom influences its bonding relationships with other atoms. Consequently, it is challenging for these diffusion models to capture the chemical dependencies between atom-atom and substructure-substructure interactions in one-shot generation. Autoregressive methods, from Pkt2Mol to FragGen, generally perform better on LBE and QED metrics by decomposing the joint distribution *p*(*A*_*a*_, *E*_*a*_|*N*_*a*_) into *p*(*E*_*a*_|*N*_*a*_) · *p*(*A*_*a*_|*N*_*a*_, *E*_*a*_), which effectively decouples the entangled relationships and compels the model to learn them. Notably, some autoregressive methods, like FragGen, show improved performance in molecular properties (like FragGen SA: 0.72 compared to Pkt2Mol SA: 0.34), however, they can only be trained on a limited number of molecules since not all molecules can be decomposed to the fragments they define, thus further reducing the learned chemical space. The other two chemical language models also show some improvements in molecular properties such as SA, but they cannot match the performance of our ECloudGen in terms of the LBE metric. In summary, molecules generated by ECloudGen not only bind tightly to target proteins but also possess rational chemical structures, thereby advancing the structure-based molecule generation models closer to real-world applications.

**Table 1.** Evaluation of ligand binding efficiency LBE and other molecular properties for baselines and ECloudGen. The standard deviation table is separately listed in SI. ↓ means the smaller the value, the better the performance, and ↑ means the opposite. Values in **bold** refer to the best metrics. AR-A/F denotes the autoregressive atom-wise/fragment-wise strategy, Chem-L denotes the chemical language strategy.

| Class | Methods | LBE (↓) | QED (↑) | SA (↑) | Lipinski (↑) | LogP(-2-5) |
| --- | --- | --- | --- | --- | --- | --- |
| Ref. | Test Set | -0.34 | 0.56 | 0.73 | 4.89 | 0.10 |
| AR-A | Pkt2Mol | **-0.42** | 0.57 | 0.34 | 4.93 | 0.76 |
| AR-A | ResGen | -0.41 | 0.55 | 0.32 | 4.95 | 1.64 |
| AR-A | GraphBP | -0.33 | 0.56 | 0.48 | 4.77 | 1.29 |
| AR-F | FLAG | -0.31 | 0.52 | 0.58 | 4.87 | 0.45 |
| AR-F | FragGen | -0.36 | 0.57 | 0.72 | 4.86 | 1.27 |
| Diff | DiffBP | -0.33 | 0.49 | 0.43 | 4.78 | 0.45 |
| Diff | DiffSBDD | -0.35 | 0.55 | 0.50 | 4.85 | 0.11 |
| Diff | TargetDiff | -0.33 | 0.57 | 0.48 | 4.78 | 0.62 |
| Chem-L | Lingo3dMol | -0.28 | 0.47 | 0.65 | 4.50 | 0.74 |
| Chem-L | PrefixMol | -0.36 | 0.51 | 0.69 | 4.83 | 0.08 |
| **Our** | **ECloudGen** | **-0.42** | **0.70** | **0.79** | **5.00** | **0.09** |

### ECloudGen Accesses a Broader Chemical Space

The first experiment demonstrated that ECloudGen achieved comprehensive improvement in both binding efficiency and molecular properties, which we attribute to its access to a broader chemical space compared to baseline models. In this study, we aim to quantitatively measure the chemical space, further affirming ECloudGen’s capability to explore a more extensive chemical space. Previous studies, such as Pkt2Mol, often used internal diversity (IntDiv) to measure chemical space; however, this widely adopted metric is not optimal. According to point-set topology, a robust measure of chemical space should satisfy subadditivity and dissimilarity preference axioms^34^, whereas IntDiv only fulfills the latter, rendering it suboptimal. Therefore, we adopted two metrics that respect both these axioms—Maximum Exclusion Circle (Circles)^34^ and Bottleneck^35^—which better measure the chemical space coverage and diversity.

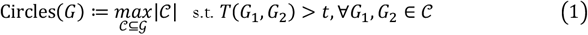

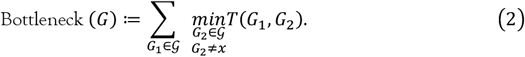

Where *G* denotes the set of generated molecules, *T* represents Tanimoto similarity. Intuitively, the ‘Circles’ metric quantifies the number of mutually exclusive circles required to cover the generated molecules within the chemical space; concurrently, the ‘Bottleneck’ metric calculates the sum of the shortest distances between each molecule and its closest neighbor. The results of the Circles and Bottleneck metrics, presented in **Figure 2A-B**, demonstrate that ECloudGen achieves the highest values for both of them. Specifically, ECloudGen scores the greatest number of mutually exclusive circles in the chemical space (85 vs second-best 63), indicating a broader coverage of chemical space. Moreover, ECloudGen exhibits the largest bottleneck (1200 vs second-best 1063), suggesting that generated molecules are widely dispersed across the chemical space, thereby indicating a structurally unique and diverse molecular set. The expansive chemical space explored by ECloudGen can be attributed to its novel approach of introducing no-structure data for training. As illustrated in **Figure 2C**, the learning objective is transferred from the traditional graph representation *p*(*G*|*p*) to our electron cloud representation *p*(*C*|*p*), where each electron cloud can accommodate several ball-and-stick molecules, thereby increasing the training data volume. In summary, ECloudGen provides a novel and efficient strategy to address the “Chemical Space Generation Paradox”, fundamentally improving the generation quality.

**Figure 2.**
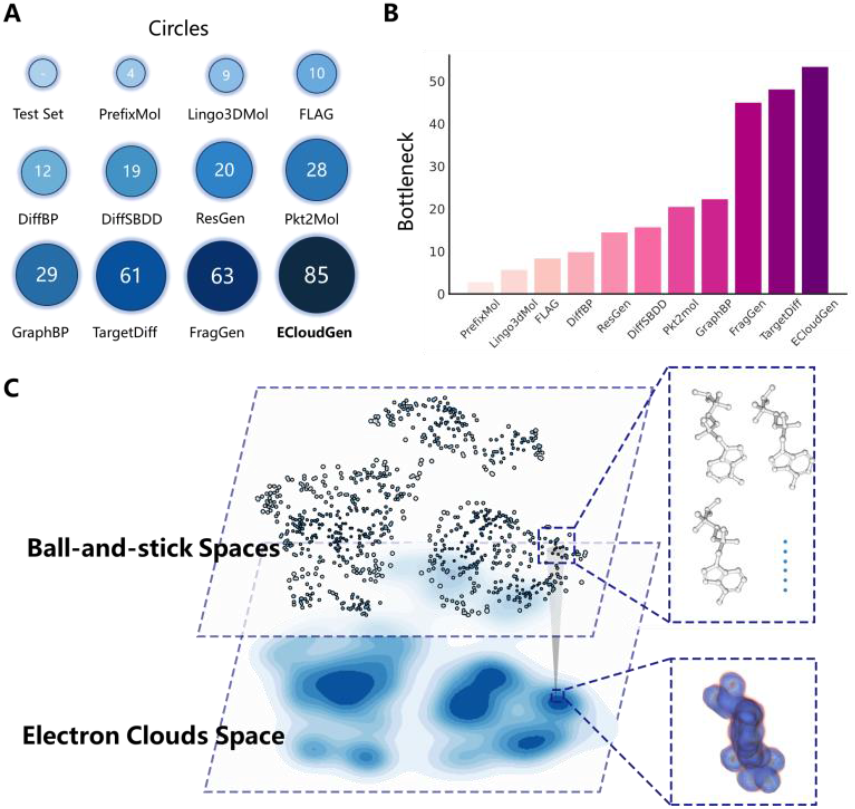
**A)** Illustration of the Maximum Exclusion Circle metric for different methods, where larger and darker circles represent higher Circles metric values. Specific numbers are put in **Table S1. B)** Scores of the Bottleneck metric for different methods, where taller and darker bars indicate higher Bottleneck metric values. **C)**. Leveraging the latent variable approach, ECloudGen maps a set of discrete molecules into a small patch in the latent space, bridging the protein-ligand and ligand-only data.

### ECloudGen Provides Model-level Intrepretability

Another major benefit of our unique approach is that our physically meaningful latent variable, electron clouds, acts as a means of interpretation, which is particularly useful in AI-decision processes requiring human intervention, especially in drug design^36^. Previous structure-based drug design approaches, like DiffBP^8^ and ResGen^5^, directly output an array of molecules from the given empty protein pockets. Though they prove successful in designing tightly binding molecules, many chemist users still remain concerned about the rationale behind the model’s generation of specific molecules within the pockets. Interpretability has always been a critical research direction for AI models. Therefore, we have constructed a pipeline in ECloudGen that incorporates model-level interpretation into the generation process, as illustrated in **Figure 3**. Initially, users obtain electron agents from the ECloud Latent Diffusion module, complete with their E-Confidence ranking scores. Subsequently, users can select electron clouds based on their interest or those with higher E-Confidence scores for subsequent generations. We demonstrate this process with a concrete example.

**Figure 3.**
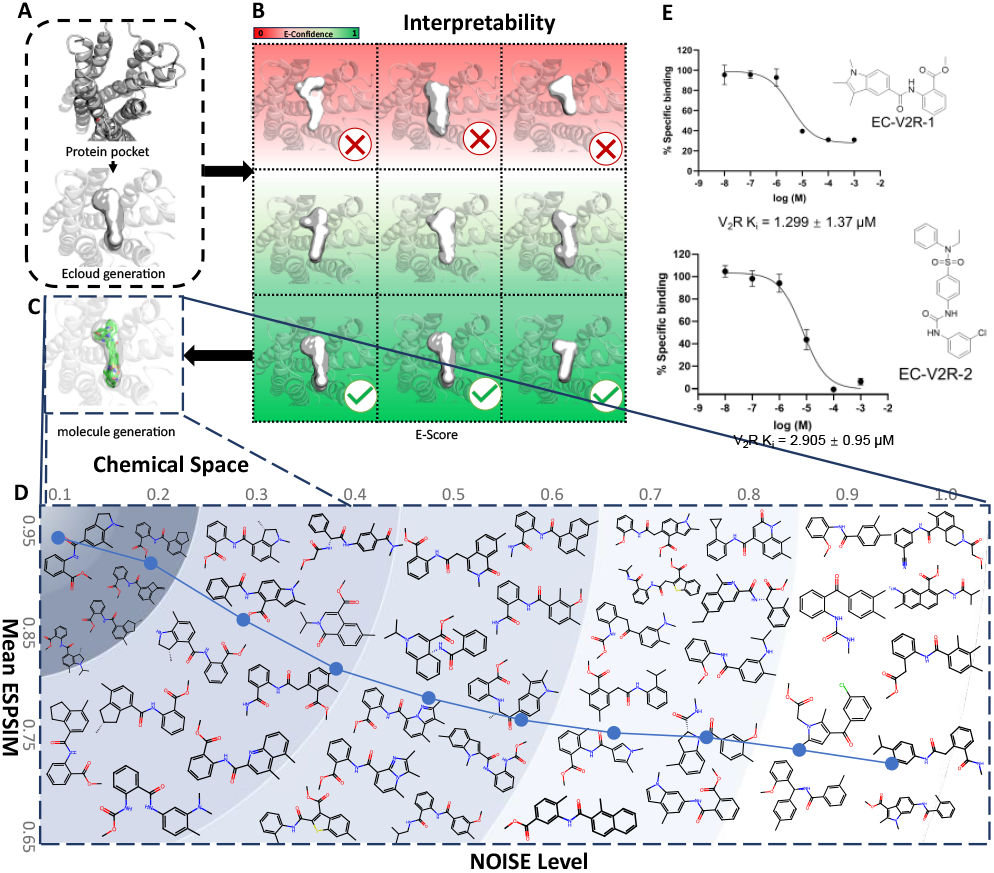
**A)** The ECloud Latent Diff Module generates electron clouds within the V2R receptor. **B)** Chemist users choose the fitting electron clouds as latent variable *C*, assisted by E-Score’s rankings. **C)**. Starting with the chosen *C*, **D)**. ECloud Decipher maps *C* onto a continuous chemical space. In this space, the further from the center, the greater the variation in molecular structures. **E)**. Two molecules were selected for synthesis and bioassay testing, yielding results with affinities in the micromolar range.

Vasopressin 2 Receptor (V2R) is a critical G-Protein-Coupled Receptor that serves as a therapeutic target for renal dysfunction^37^. Antagonist drugs such as Tolvaptan^38^ are developed to target this receptor, but exhibit serious side effects^39^. This case is selected to demonstrate how to employ the interpretable generation protocol with ECloudGen for lead compunds design. Specifically, ECloudGen initially generated a series of electron cloud agents from the V2R pocket, as shown in **Figure 3A**. We then selected three electron clouds with E-Confidence scores above 0.8 (**Figure 3B**) to complete the molecule generation (**Figure 3C**), resulting in the structures shown in **Figure 3D**. These generated molecules exhibit electron similarity to their corresponding intermediate electron clouds, demonstrating the effectiveness of using electron cloud intermediates as a means for model generation interpretation. We conducted a rapid validation of the generated results from two electron clouds of different sizes, and obtained two molecules with *μ* level activity (**Figure 3E**). This naïve proof of concept expermeint demonstrates how chemists could discard or select molecules from their attribution to the electron clouds generative agents, showing a unique aspect of ECloudGen.

### ECloudGen Enables 3D Protein-specific Single-objective Molecular Optimization

In previous experiments, we demonstrated that compared to current models, ECloudGen can efficiently design ligands for target proteins that not only exhibit high efficacy and superior properties but also explore a broader and more continuous chemical space. Thanks to our latent variable appraoch, ECloudGen also possesses optimization abilities that enable molecular optimization toward desired target properties. Existing structure-based molecular generation studies often overlook the importance of molecular optimization, typically relying on the posteriori approach^40^: generating a large number of molecules followed by filtering based on desired properties. However, this approach requires the model to generate molecules that are both diverse and pharmaceutically feasible, posing significant challenges to current models. To address this, we designed two molecular optimization strategies within ECloudGen: the meta-dynamics-inspired^41,42^ ESpaceEvo module and the PSO-based^43^ EPSO module. The former is model-specific, requiring derivative information of the objective function, while the latter is model-agnostic and only needs the objective function values. The advantage of the former lies in providing a complete molecular optimization trajectory, aiding chemists in tracking and interpreting the model’s decision-making process; the latter’s benefit is its ease of use, as it operates without the need for constructing a surrogate model on the ECloudGen chemical space. Therefore, our analysis here focuses on the EPSO module. For more information regarding ESpaceEvo, interested readers are referred to **S3**.

Given the lack of baselines for structure-based molecular optimization, we compared our model against those without protein pocket constraints. In this setting, the protein condition input for ECloud Decipher is set to zero paddings. Following the Modof^46^ methodology, QED is chosen as the test objective function, and 800 test molecules within a QED range of [0.7, 0.8] serving as starting points. Four models serve as baselines: JTNN^44^, HierG2G^45^, Modof-pipe, and Modof-pipe(m)^46^. JTNN and HierG2G optimize the input molecular graph using tree and fragment graph structures, respectively, through variational inference. Modof-pipe modifies individual fragments iteratively, whereas Modof-pipe(m) utilizes beam search to select the top-5 candidates in each iteration. As shown in **Table 2**, ECloudGen outperforms all baselines under two definitions of optimization success. For instance, it achieves a success rate of 79.20% for QED >0.9, a 5% improvement over the second-ranked HierG2G, and 89.20% for QED Improvement >0.1, approximately a 2% improvement over the second-ranked Modof-pipe(m). It is noteworthy that JTNN and HierG2G are model-specific, necessitating additional training of surrogate functions in the latent space, which inevitably introduces additional errors. Both Modof variants require obtaining the training pairs from Low QED to High QED molecules with single fragment modifications, making them difficult to generalize to new properties of interest. All these methods struggle to scale effectively when faced with more complex real-world objective functions. In contrast, ECloudGen, through its EPSO module, achieves plug-and-play functionality without the need for additional models or data preparation. We will demonstrate its versatility in the next section.

**Table 2.** Single objective optimization (QED) results.

| Methods | QED > 0.9 |  |  | QED Imprv > 0.1 |  |  |
| --- | --- | --- | --- | --- | --- | --- |
|  | Rate (%) | Imprv | Sim | Rate (%) | Imprv | Sim |
| JTNN | 60.50 | 0.17±0.03 | 0.47±0.06 | 67.38 | 0.17±0.03 | 0.47±0.07 |
| HierG2G | 75.12 | 0.18±0.03 | 0.46±0.06 | 82.38 | 0.17±0.03 | 0.46±0.06 |
| Modof-pipe | 40.00 | 0.17±0.03 | 0.54±0.08 | 70.00 | 0.16±0.03 | 0.51±0.08 |
| Modof-pipe(m) | 66.25 | 0.18±0.03 | 0.48±0.07 | 87.62 | 0.17±0.03 | 0.48±0.07 |
| ECloudGen (ours) | **79.20** | **0.18±0.03** | 0.47±0.06 | **89.20** | **0.18±0.03** | 0.47±0.06 |

### ECloudGen Enables Multi-objective Goal-directed Molecular Optimization

We have demonstrated the single-target optimization capabilities of ECloudGen. In this section, we expand ECloudGen’s optimization to multiple objectives, aligning more closely with the real-world drug design scenario. Similar to single-target optimization, the challenge of multi-target molecular optimization based on protein pockets remains unresolved. Thus we aim to illustrate the ECloudGen’s ability in multi-objective optimization with a case study rather than a comprehensive model comparison. Additionally, most multi-target molecular optimization methods are model-specific, necessitating the training of surrogate prediction models and generally restricted to continuous objective functions. However, molecular optimization in real-world scenarios often involves abstract, discrete objective functions, such as favoring molecules with specific functional groups, which render many model-specific models ineffective. In contrast, ECloudGen mitigates these limitations through its model-agnostic and multi-objective EPSO module. We use the redesign of endogenous ligands to exogenous inhibitors for Bromodomain containing protein 4 (BRD4) as a case study to demonstrate the complexity of real-world objective functions and the applicability of ECloudGen’s model-agnostic optimizer in such contexts.

BRD4 belongs to the bromodomain and extra-terminal domain family, whose overexpression is closely associated with the development of various cancers^47^. BRD4 functions by recognizing acetylated lysine residues in histones H3 and H4. Thus, a natural approach to drug design against BRD4 is to design exogenous inhibitors that compete with acetylated lysine, thereby influencing BRD4’s biological functions in transcription and exerting therapeutic effects^48^. In this scenario, the goal is to design an exogenous ligand that maintains the endogenous ligand’s binding pattern with BRD4 protein. We established a design pipeline using ECloudGen, starting with the electron cloud of the “endogenous ligand” acetylated lysine, as shown in **Figure 4A**. In selecting objective functions, MolSearch^49^ found that over-optimization of molecular properties such as QED and SA results in smaller molecules; therefore, we reframed molecular property optimization as a constrained problem. Following the strategy in PocketFlow^15^ for screening AI-generated molecules, we defined the molecular property optimization function as (1) QED>0.6, (2) SA>0.6, (3) molecular weight greater than 250, and (4) Lipinski=5. Furthermore, the co-crystal complex structure of BRD4 with histones^50^ points out that acetylated lysine forms key hydrogen bonds with Asn140 and Tyr97 of BRD4. To accommodate these crucial interactions, we we set an additional optimization objective of (5) H_acceptor ≥ 1 for the molecules.

**Figure 4.**
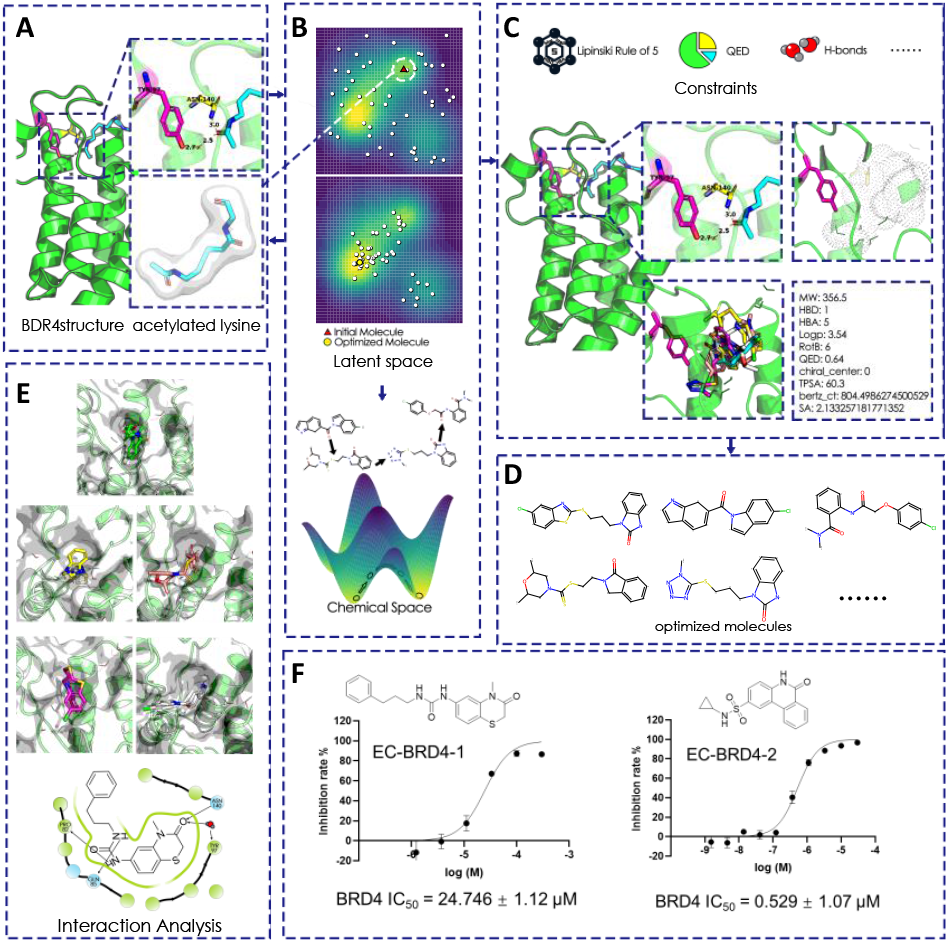
**A)** BRD4 functions by binding with “endogenous ligand” acetylated lysine in histones H3 and H4. **B)**. EPSO module perform molecular optimization on the latent space. **C)**. Five objectives of BRD4 redesign of endogenous ligand, three for molecular properties and two for special consideration in BRD4 case. **D)**. ECloudGen optimized molecules. **E)**. Docking positions of several oprimized molecules with BRD4 targets. **F)**. Two molecules were selected for synthesis and bioassay testing, yielding results with affinities in the micromolar range.

After defining the optimization objectives, as illstrated in **Figure 4C**, we used ECloudGen to perform multi-objective molecular optimization starting from the “endogenous ligand”, acetylated lysine, as depicted in **Figure 4B**. The statistical metrics of the optimized molecules are displayed in the **Table S2**. After 50 PSO evolutions, almost all optimized molecules satisfied the five optimization constraints, as shown in **Figure 4D**. To further narrow down the selection space, we docked molecules into the BRD4 proteins and ranked them using docking scores. It can be observed that the optimized molecules shown in **Figure 4E** remain the key interactions with BRD4 binding pockets, implying the validity of our proposed optimization objectives. Finally, two molecules were selected by our medicinal chemists for synthesis and bioassay, yielding two molecules with activities of 25μMol and 0.5μMol, as shown in **Figure 4F**. This redesign of the endogenous ligand showcases how ECloudGen emulates the thinking of a medicinal chemist and provides a concrete reference for its application in multi-objective molecular optimization campaigns.

## Conclusion

In this work, we first identify one of the biggest challenges in structure-based molecular generation, naming it as the Sparse Chemical Space Generation Paradox. To address this, we propose introducing a latent variable *C*, which decomposes *p*(*G*|*p*) = *p*(*G*|*p, C*)*p*(*C*|*p*), thereby bridging the limited protein-ligand paris and ligand-only data. Inspired by quantum physics and modern generative models, we define electron clouds as the informative latent variable and develop a model named ECloudGen based on this concept. Through our comprehensive analysis, including both benchmarking models and real-world drug design projects, we conclude that ECloudGen can generate molecules that bind efficiently and exhibit enhanced molecular properties, showing a 19% improvement in QED and a 9.7% improvement in SA compared to the second-best baseline, respectively. Furthermore, we also demonstrated the model-level interpretability afforded by the electron clouds, as exemplified in the V2R drug design project, where we successfully obtained two lead molecules with micromolar affinity. By utilizing CEMP, the introduced contrastive learning technique, we have structured ECloudGen’s chemical space in an ordered manner and built a model-agnostic EPSO optimization module on it. The efficacy of EPSO was showcased through a single-objective oracle experiment and a five-objective constraint redesign of the endogenous ligand for BRD4. In the future, we anticipate broader applications of ECloudGen in drug design campaigns. Its unique methodology could also be further exploited in tasks such as molecular property prediction and chemical synthesis, where the underlying physical information encapsulated by electron clouds can facilitate deeper learning of fundamental knowledge.

## Methods

### Problem Statement

As discussed in the Introduction, the data scarcity issue is severe in structure-based molecular generation. We analyze this issue from the perspective of generative models and have termed it the Sparse Chemical Space Generation Paradox. To address this, we propose introducing a latent variable that bridges protein-ligand pairs and ligand-only data, thereby granting the model access to a broader chemical space, comparable at least to that of existing chemical libraries. The rationale for this proposal is grounded in probability decomposition, expressed as *p*(*G*|*p*) = *p*(*G*|*p, C*)*p*(*C*|*p*), where *G* is molecules, *p* is proteins and the *C* is desired latent variable. Inspired by quantum physics, we have identified electron clouds, denoted as *ρ*, as the latent variable *C*, due to its informativeness and ability to introduce interpretability and controllability into the generation process. In this work, we use the grid data structure to represent the electron clouds and formulate the problem as follows: *p*(*C*|*p*) aims to generate the latent variable 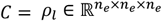, given protein *p*, where *n*_*e*_ is the number of grids. *p*(*G*|*p, C*) aims to generate molecules *G*, given the latent variable *C* = *ρ*_*l*_ and protein *p* . In the context of conditional generation, with the optimization goal of 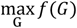 subject to constraints 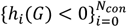, ECloudGen seeks to generate a diverse set of molecules 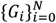 that respect the optimization objectives.

### Model Architecture

#### ECloud Latent Diffusion

The first part of ECloudGen is to learn *p*(*C*|*p*), i.e., generating electron clouds *ρ*_*l*_ = *C* from specified protein pockets. We formulate it as a conditional 3D generation problem. Inspired by the success of diffusion models in image generation^6,22,51^, we leverage latent diffusion to generate high-fidelity electron clouds given to proteins. To achieve this, we developed the ECloud Latent Diffusion module, an adaptation of the diffusion model that operates on latent space. Compared to vanilla diffusion, our module is not only memory-efficient but also adept at capturing complex geometric patterns in latent space, thereby boosting both the efficiency and the efficacy of the generation process.

#### Protein Conditional AutoEncoder (P-CAE)

To construct the latent space for the diffusion process, we begin by training a protein-conditional autoencoder. In this setup, the protein *ρ*_*p*_ conditions the mapping of the ligand electron cloud *ρ*_*l*_ into the compact latent space:

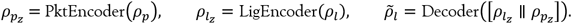

Here, 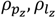 denote the embeddings of *ρ*_*p*_, *ρ*_*l*_, and 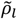 is the reconstructed ligand electron cloud. The PktEncoder, LigEncoder, and Decoderare are essentially 3D CNN models with both sub-sampling and up-sampling techniques to facilitate the encoding and decoding processes.

#### Conditional Latent Diffusion

In contrast to vanilla diffusion, which operates directly in the data space, our approach applies noising and denoising processes within the P-CAE encoded latent space. During the noising process, Gaussian noise with variances {*β*_0_, …, *β*_*T*_} is added to 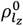 within the latent space, where the posterior distribution *q* can be represented as:

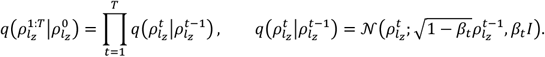

From the equations above, the closed-form expression for the probability distribution at any time step *t* in the noise-adding process is:

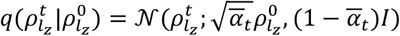

where *α*_*t*_ = 1 − *β*_*t*_ and 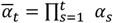. This deterministic noise addition process aims to evolve the original molecular electron cloud 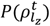 into a Gaussian distribution 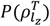. To generate a new electron cloud 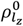 within the latent space, we first sample from the noise distribution 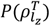. Then, we condition on 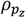 and reverse the noising process, transforming the sampled noise back into an actual electron cloud 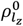. The denoising process can be expressed as:

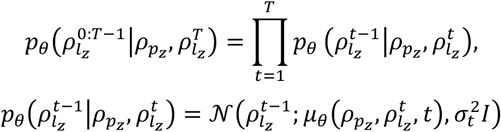

Here, *p*_*θ*_ represents the distribution with parameters to be estimated, *μ*_*θ*_ is the neural network estimating the mean of the *p*_*θ*_ and 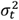 is a user-defined variance. Equation (8) illustrates how 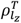 evolves step-by-step through 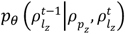 into the initial distribution sample 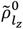. After obtaining 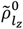, we use the P-CAE’s decoder to reconstruct the electron cloud from the latent space:

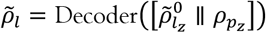

#### Training Objective

The training objectives for optimizing the 3D Conditional Latent Diffusion module can be summarized as follows. First, we consider the auto-encoder loss associated with compressing molecular electron clouds:

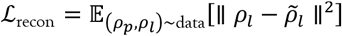

where *ρ*_*l*_ and 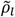 represent the input and reconstructed ligand electron clouds, respectively. Second, we consider the pocket-conditional latent diffusion loss which can be expressed as:

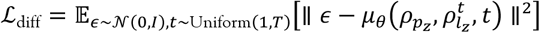

Here, ℒ_diff_ denotes the standard diffusion loss, instructing the model on the denoising of electron clouds during generation. Furthermore, we consider two regularization losses, ℒ_*t*=0_ and ℒ_*t*=*T*_, to modulate the diffusion process at starting *t* = 0 and termination *t* = *T* steps, enhancing stability and sampling quality:

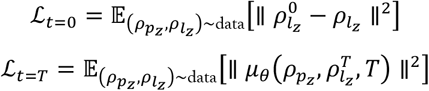

The goal of ℒ_*t*=0_ is to ensure the model’s initial output closely aligns with the original latent representation, while ℒ_*t*=*T*_ aims to control the noise introduction, preventing excessive magnitudes in the latent space encoder output. Additionally, we have a clash loss ℒ_clash_ to augment protein-awareness and mitigate steric clashes:

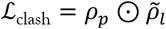

Here, *ρ*_*p*_ denotes the input protein electron cloud, 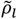 is the reconstructed electron clouds, and ⊙ is the element-wise multiplication.

#### ECloud Decipher

The second part of ECloudGen is to learn *p*(*G*|*p, C*), aiming to utilize the protein and latent variable information to generate the corresponding molecules, which we formulate as a 3D object captioning problem. To achieve this, we designed the ECloud Decipher module, which leverages a Llama-like^7^ backbone and 3D-Unet^52^ to make the model conditional on proteins and the given latent variable. In addition, we introduce the Contrastive ECloud-Molecule Pre-training (CEMP) strategy to structure the chemical space, enabling subsequent construction of the model-agnostic optimization module.

#### Llama-like Backbone

Large language models have been proven effective in natural language processing^7,53^. Therefore, we want to leverage its powerful architecture to interpret 3D structural information of electron clouds and topological information of chemical structures. Specifically, we adopt the RoFormer^54^, a rotary position embedding-enhanced transformer architecture used in Llama^7^, as the ECloud Decipher backbone. RoFormer’s core feature, Rotational Position Encoding (RoPE), integrates absolute positional encodings at the input stage and calculates relative positional encodings between inputs during attention operations. To simplify, we present the RoFormer layer without multi-head dimensions:

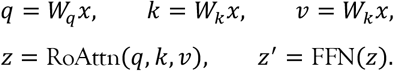

Here, *W*_*q*_, *W*_*k*_, *W*_*v*_ are the weight matrices query, key, and value, respectively. RoAttn denotes the rotational attention computation module, and FFN is the ffully connectednetwork. The expansion of the RoAttn module can be written as follows:

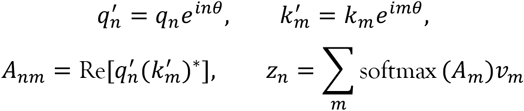

Here, 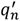 and 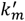 are the complex features of the *n*-th query and the *m*-th key, respectively. *θ* denotes a predefined base angle, *d*_*k*_ is the embedding dimension, and Re represents the real-part operation. After the softmax on the attention map *A, z*_*n*_ is obtained as the output feature.

#### ECloud Caption Loss

The input of the second part of ECloudGen includes the latent variable *C*, encapsulating molecular information encoded by electron clouds, and the protein *p*, aiding *C* in generating corresponding molecules. We use the 3D-Unet backbone to encode the electron clouds into the chemical space, followed by decoding them into the corresponding molecules using RoFormer_d_. Protein conditions are integrated using a cross-attention mechanism within RoFormer_d_ . If no corresponding proteins are available, a virtual protein vector *b* serves as an alternative. The training objective is defined as:

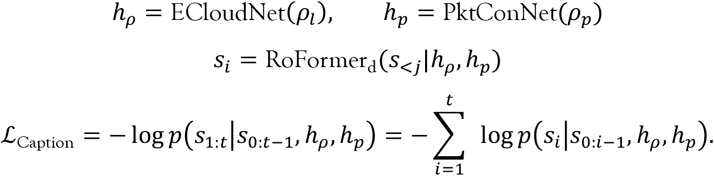

where *h*_*ρ*_, *h*_*p*_ represent the latent chemical features of the electron cloud *ρ*_*l*_ and protein *ρ*_*p*_, *s*_*i*_ is the *i*-th token of molecules, RoFormer_d_ denotes the RoFormer acting as the decoder, and ℒ_Caption_ is in the form of the negative log-likelihood loss function.

#### CEMP for Structurally Ordered Space

Molecular optimization is one of the most important elements in drug discovery, including the optimization of the binding affinity to enhance drug efficacy, as well as the improvement of ADMET profiles to enhance safety and bioavailability^55,56^. The current structure-based molecular generation models often grapple with the limited size of accessible chemical spaces, while ECloudGen addresses this by tapping into a broader chemical space. To fully utilize this advantage, we want to make chemical space structurally ordered, which means that similar molecules are grouped together in the learned chemical space, facilitating the smoothing optimization in ECloudGen. A common approach is to employ a variational autoencoder (VAE) architecture^57^, which naturally clusters data in latent space through its Gaussian prior constraint. However, the simplicity of the Gaussian constraint makes it inefficient to capture the complexity of the chemical space. One possible modification is to relax this constraint through the adoption of a mixture of Gaussian distributions^58^; nevertheless, this still does not solve the problem fundamentally.

To overcome the above limitation, we then explored another approach—contrastive learning. The basic idea behind contrastive learning is to bring positive samples closer in the representation space while pushing negative samples away. Although it is primarily used in the pre-training to improve performance in downstream tasks^59,60^—seen in Contrastive Language-Image Pre-training (CLIP)^61^, which achieves superior zero-shot classification by embedding images and text into a shared space—its principles are found to be ideally suited for our purpose. Therefore, we proposed Contrastive ECloud-Molecule Pre-training (CEMP) to structure the complex chemical space of ECloudGen, thus facilitating the extension of our generative model to include molecular optimization tasks. To achieve this, we begin by writing down the features of contrastive elements:

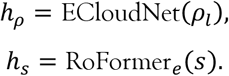

Here, *h*_*ρ*_ and *h*_*s*_ denote the respective embeddings for electron cloud and molecule in the chemical space. The electron cloud of a given molecule is considered a positive sample, whereas electron clouds from different molecules are considered negative samples. To perform contrastive learning on these samples, the InfoNCE loss^62^ can be used:

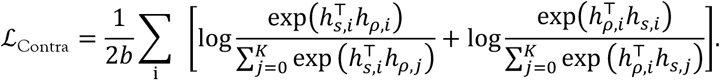

where ℒ_Contra_ aims to maximize the similarity between positive pairs while minimizing the similarity between negative pairs. CEMP facilitates flexible clustering in the latent space, helping the model distill hidden physicochemical knowledge in a self-supervised way.

#### Model-agnostic Molecular Optimization

As previously discussed, ECloudGen, through its latent variable approach combined with ecloud-molecule contrastive learning, possesses a structurally ordered chemical space. This property ensures that molecular structural similarity correlates with spatial proximity, thereby facilitating continuous molecular optimization. In this context, our aim is to develop a model-agnostic optimizer operating within this learned space. The term “model-agnostic” implies that detailed information about surrogate models (objective functions) is not required. For example, gradient evolution on the learned space is not a model-agnostic method as it needs the gradient of objective functions, meaning we need to additionally train a model *g* to find the map from molecules *m* to conditions *h*, i.e., *h* = *h*(*g*(*m*)). Therefore, a model-agnostic optimizer holds great value in simplifying the molecular optimization process and enhancing the model’s user-friendliness.

In practical terms, we utilize particle swarm optimization to construct the ECloudGen-PSO (EPSO) module. Initially, we place *n* particles on the space to represent *n* optimized molecules, with the optimization process evolving over *t* steps as described below:

Velocity update

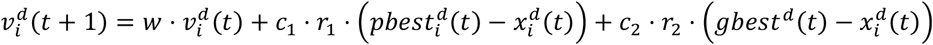

where *d* is the *d*-th dimension, 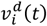 is the velocity of the *i*-th molecule, 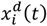 is its position of the learned space, 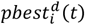 and *gbest*^*d*^(*t*) are historical best solutions of this and all optimized molecules, *w* is the inertia weight, *c*_1_, *c*_2_ are learning factors, and *r*_1_, *r*_2_ are random numbers in the interval [0,1]. Once obtained the velocity of molecules, we update their positions and evaluate their optimizing fitness:

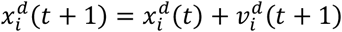

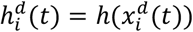

Based on the evaluated fitness, we update the local best position and the global best position as follows:

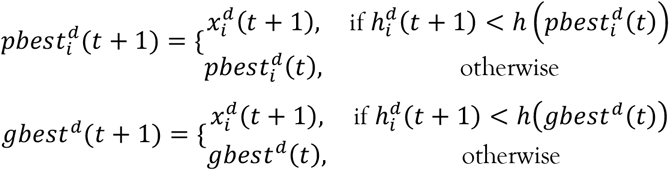

After *T* evolutionary steps, the final optimized molecules are obtained from the global best positions *gbest*.

#### Electron Clouds Computation

Electron clouds, also known as electron density, are the foundation of quantum chemistry. As discussed in the Introduction, most molecular systems’ physicochemical properties are electron clouds’ functions. Additionally, electron clouds unify all interatomic forces within and between molecules, including bonded and non-bonded interactions^20^. In this regard, electron clouds provide a more informative representation compared to the canonical ball-and-stick representation. Essentially, electron clouds can be obtained by solving the Schrödinger equation^63^ as follows:

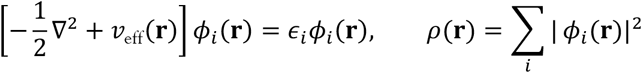

Here, *ϕ*_*i*_(**r**) refers to the electron’s wave function, *ϵ*_*i*_ denotes the energy eigenvalue, 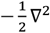 represents the kinetic energy operator, and *v*_eff_(**r**) is the effective potential. *ρ*(**r**) refers to the electron cloud, indicating the probability density of the presence of an electron at a specific position **r**. However, directly solving the Schrödinger equation to obtain electron clouds is computationally expensive. To make the problem tractable, an efficient semi-empirical GFN2-xTB method^64^ is utilized to calculate electron clouds. GFN2-xTB is an adaptation of the tight-binding approach, which is specifically designed for chemical systems and has been validated as sufficiently accurate for calculating electron clouds.

### Molecular Properties Decription

#### Quantitative Estimate of Drug-likeness (QED)

As described in its development paper^27^, QED is a metric that provides a quantitative estimate of drug-likeness based on a molecule’s desirability across a range of properties. It is a composite score derived from molecular properties that are typically considered favorable for drug candidates.

#### Synthetic Accessibility (SA)

The Synthetic Accessibility Score, as outlined in Ref^28^, measures the ease with which a molecule can be synthesized. The calculation of the SA score is based on several factors, including molecular complexity (like containing unusual rings) and fragment contributions.

#### The partition coefficient between octanol and water (LogP)

Logp is a measure of a molecule’s hydrophobicity^30^. It is crucial in determining a compound’s absorption and distribution characteristics, influencing its pharmacokinetic profile. Generally, it is considered that a LogP value between -2 and 5 represents a reasonable range for potential drug candidates.

#### Lipinski’s Rule of Five (Linpinski)

Introduced in its original development^29^, Lipinski’s Rule of Five predicts the drug-likeness of compounds based on four key physical properties: no more than 5 hydrogen bond donors, no more than 10 hydrogen bond acceptors, a molecular weight under 500 Dalton, and an octanol-water partition coefficient (LogP) not greater than 5. These rules are used to evaluate the likelihood of success in the later stages of drug development.

#### Ligand Binding Efficiency (LBE)

To address the inherent bias of the Vina score favoring larger molecules due to its calculation method — summing up the individual atom or fragment energy contributions — we adopt the Ligand Binding Efficiency (LBE). LBE normalizes the Vina score by the number of atoms *N*_*a*_ in the molecule, calculated as 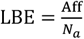.

## Supporting information

Supp All

## Data and Code Availability

The data and source code of this study is freely available at GitHub (https://github.com/HaotianZhangAI4Science/ECloudGen) to allow replication of the results.

## Supporting Information

**Part S1**. Potential Benefits of Learning on Electron Clouds; **Part S2**. Metric Values of Chemical Space Measurement; **Part S3**. Model-specific Optimizer of ECloudGen: ESpaceEvo; **Part S4**. Illustration of Generated Molecules; **Part S5**. Chemical Synthesis; **Part S6**. Bioassay Methods **Table S1**. Chemical space measurement metrics (Circles and BN) of different models. **Algorithm S1** ESpaceEvo Method. **Figure S1** Illustration of metadynamics-inspired ESpaceEvo. **Figure S2** Generated examples from different methods.

## Acknowledgments

This study was supported by the National Key Research and Development Program of China (2021YFE0206400), and the National Natural Science Foundation of China (22220102001, 82204279, 92370130).

