## Supplementary material for "ECloudGen: Leveraging Electron Clouds as a Latent Variable to Scale Up Structure-based Molecular Design": Supp All

### S1 Potential Benefits of Learning on Electron Clouds

Although the ball-and-stick model is an early and very familiar concept in chemistry, it is important to note that it is only an approximation based upon classical mechanics[1]. The electron cloud model, on the other hand, is a quantum-based, foundational representation, as it describes the probability density of electrons[2]. This is more in tune with the dynamical and probabilistic nature of atoms[3,4]. Learning using electron clouds offers two potential advantages:

(1) Firstly, from a physics perspective, all the intermolecular forces reduce to only one force, i.e., electromagnetic force. Examples include hydrogen bonds, van der Waals forces, and $\Pi$-$\Pi$ stacking, which are basically variants of dipole-dipole interactions[5]. By using an electron cloud-based learning model, the model bypasses the redundant classical approximation and directly imbibes the unified laws of interaction for superior generalization capabilities.

(2) Secondly, compared with learning on discrete distribution, learning on continuous fields is an easier task. In the ball-and-stick representation, the difficulty of training is from both the atom positions, which discretize the spatial domain with existence and non-existence, and atom element types, which are treated as discrete variables. In contrast, electron clouds are continuous in the spatial domain, enabling efficient utilization of the valuable protein-ligand data. Moreover, electron clouds provide a denser learning space. As shown in **Figure 2C**, the molecules which appear distinct in the chemical space tend to merge in electron cloud space, thus providing a more generalization capability of the model toward new, unseen molecules, which is a very important advantage, considering the scarcity of 3D data. Based on these intuitions, we believe that electron clouds provide a unified perspective of molecular learning, as well as a more favorable way for AI model training.

### S2 Metric Values of Chemical Space Measurement

**Table S1.** Chemical space measurement metrics (Circles and BN) of different models.

| | Pkt2Mol | ResGen | GraphBP | FLAG | FragGen | DiffBP |
|---|---|---|---|---|---|---|
| Circles | 28 | 20 | 29 | 10 | 63 | 12 |
| BN | 20.5 | 14.5 | 22.3 | 8.4 | 45.0 | 9.9 |

| | DiffSBDD | TargetDiff | Lingo3dMol | Prefixmol | ECloudGen |
|---|---|---|---|---|---|
| Circles | 19 | 61 | 9 | 4 | **85** |
| BN | 15.7 | 48.1 | 5.7 | 2.8 | **53.4** |

### S3 Model-specific Optimizer of ECloudGen: ESpaceEvo

In addition to the model-agnostic module EPSO, we also implemented a meta-dynamics-inspired optimization module, named ESpaceEvo. Specifically, gradient-based optimization is utilized to optimize molecular structures in the chemical space. To obtain diverse optimized molecules, the meta-dynamics methodology[6-8] is leveraged to bump the model out of local minima when structural changes are minimal, where the adding potential is defined as (for simplicity, we describe this in 2D):

$$\phi(H, c_i, r) = H \cdot \sum_{c_i \in \{c\}} \left( -\frac{1}{2r}(h_\rho - c_i)^2 + \frac{1}{2}(\log(2\pi) + \log(r)) \right)$$

Here, $\phi(H, c_i, r)$ is the Gaussian potential function to lift the optimization target out of local minima. $c_i$ is the position of the Gaussian potential, $r$ is its variance, and $H$ is its height coefficient. The detailed algorithm for ESpaceEvo is presented as follows:

---

**Algorithm S1** ESpaceEvo Method

---

**Input**: Initial molecule and its embedding vector $m_0$, $v_0$, objective function $h(v)$, constraint functions $\{h_i(v)\}_{i=1}^{n}$ and the corresponding Lagrange multipliers $\{\lambda_i\}_{i=1}^{n}$, optimization steps $n$, and the decoder of ECloud Decipher Decoder, the Gaussian potential $\phi(H, c_i, r)$ with the given $H$ and $r$

**for** $i = 0$ to $n_{steps}$ **do**:

    Calculate objective function $f(v_i)$

    Aggregate constraints and compute total constraint penalty $\sum_i \lambda_i h_i(v)$

    **if** $\frac{v_i \cdot v_{i-1}}{\|v_i\| \|v_{i-1}\|} < \delta$ and $\boldsymbol{i} > 0$ **then**

        Calculate adding potential $\phi(H, c_i, r)$

    **else**

        $\phi(H, c_i, r) = 0$

    **end if**

    Calculate the Lagrangian function $\mathcal{L} = f(v_i) + \sum_i \lambda_i h_i(v) + \phi(H, v_i, r)$

    Calculate the gradient $\nabla\mathcal{L}$

    Update $v_{i+1} = v_i + \nabla\mathcal{L}$

    Decode the vector $m_i$ = Decoder$(v_i)$

**end for**

**Output:** $\{m_i\}_{i=0}^{n_{steps}}$

---

EPSO implemented molecular optimization by performing gradient-based evolution directly on to-be-optimized molecules within the chemical space. Specifically, given a target, we first construct a prediction model for the given pharmacological property of interest in the chemical space. Starting with some seed molecules, we guide their evolution within the chemical space using the gradients of the prediction model to optimize the molecules. Additionally, the metadynamics-inspired approach is utilized to enable molecules to escape from basins when they become trapped in local minima, as shown in **Figure S A**.

### Single-objective Optimization with ESpaceEvo

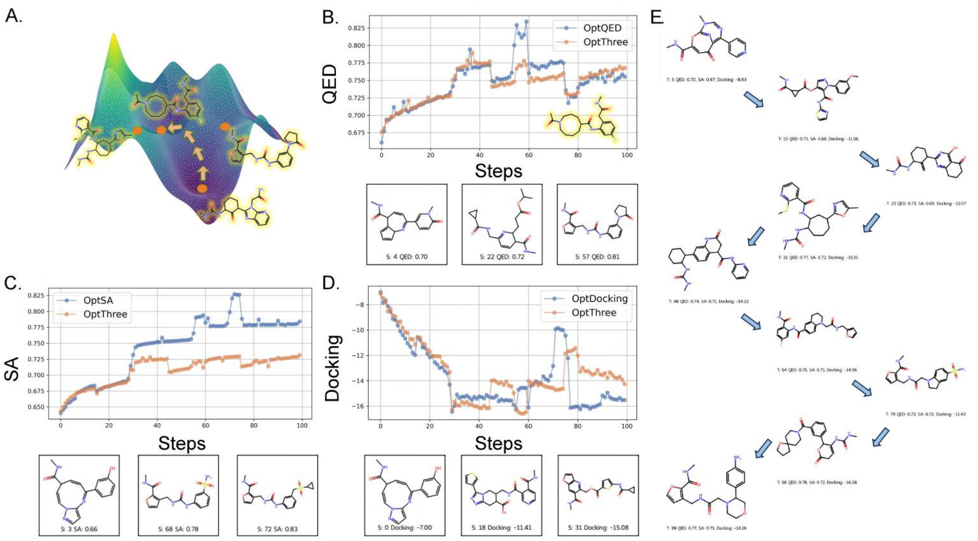


**Figure S1** A). Illustration of metadynamics-inspired ESpaceEvo. B, C, D). Objective function values for single- and multi-objective optimization, with each canvas plotting objective values under the specific single (blue) and multi-objective (orange) settings. E). Illustrated optimization trajectories of the multi-objective case.

For demonstrating the ESpaceEvo module, we select the protein with PBD ID of 4bel as our case study, optimizing for two molecular properties, QED and SA, in addition to binding affinity estimation via docking. We first conduct single-objective optimization (see illustration of blue lines in **Figure S1 B-D**. The sudden jumps observed during convergence are due to the bumping potential (meta-dynamics) applied by ECloudGen when the molecules become trapped in valleys of the chemical space. Although the optimization objective may temporarily worsen after these jumps, the molecules rapidly evolve to better values in new regions.

### Multi-objective Optimization with ESpaceEvo

In addition to the single-objective optimization, we also perform multi-objective optimization across three objectives once, as shown in **Figure S1 B-D**. We can see that the optimization outcomes for Docking and QED in multi-objective optimization closely align with those observed in single-objective optimization. However, the performance in optimizing for SA is slightly less effective, which may be attributed to the inherently lower SA values of molecules in the newly accessed regions. The illustration of trajectories for multi-objective optimization is shown in **Figure S1 E**.

### S4 Illustration of Generated Molecules

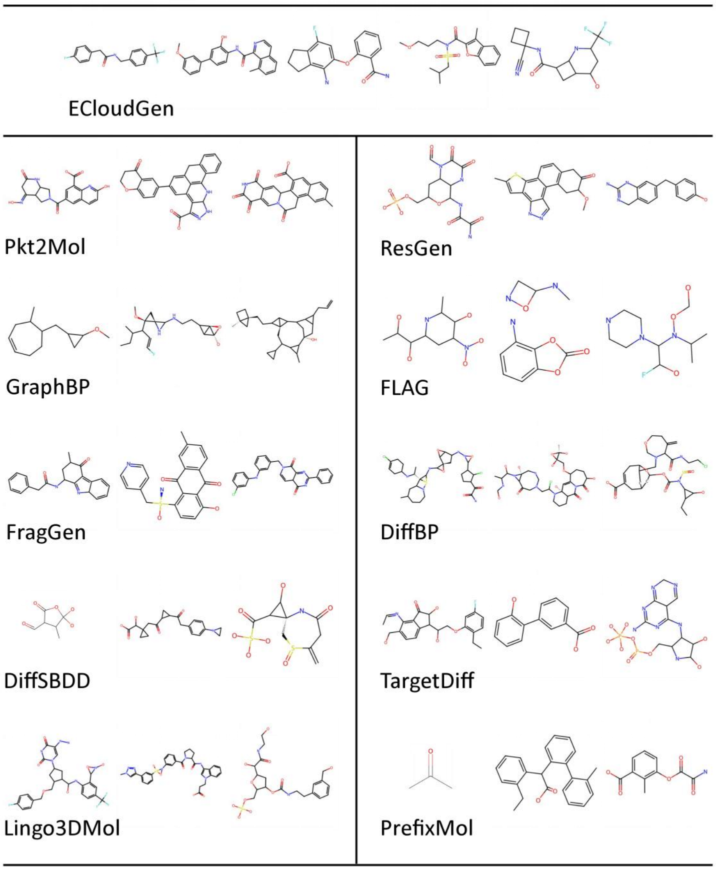


**Figure S2** Generated examples from different methods.

### S5 Chemical Synthesis

All chemical reagents and solvents were purchased from commercial sources and used without purification. Thin-layer chromatography (TLC) was carried out using silica gel plates coated with fluorescence F-254. Product spots were visualized under UV light ( λ = 245 nm or 365 nm) and stained with potassium permanganate or phosphomolybdic acid. Normal-phase silica gel column chromatography was carried out to purify crude products. $^{1}$H NMR and $^{13}$C NMR data were obtained at 400 MHz and 101 MHz, respectively, using a Bruker AV-400 spectrometer in the deuterated solvent specified. Chemical shifts are reported as δ values (parts per million) relative to the residual nondeuterated solvent signal as an internal reference. Coupling constants (*J*) are reported in Hertz. The multiplicity was defined as singlet (s), doublet (d), triplet (t), broad (br), or multiplet (m).

**Synthesis of EC-BRD4-1**

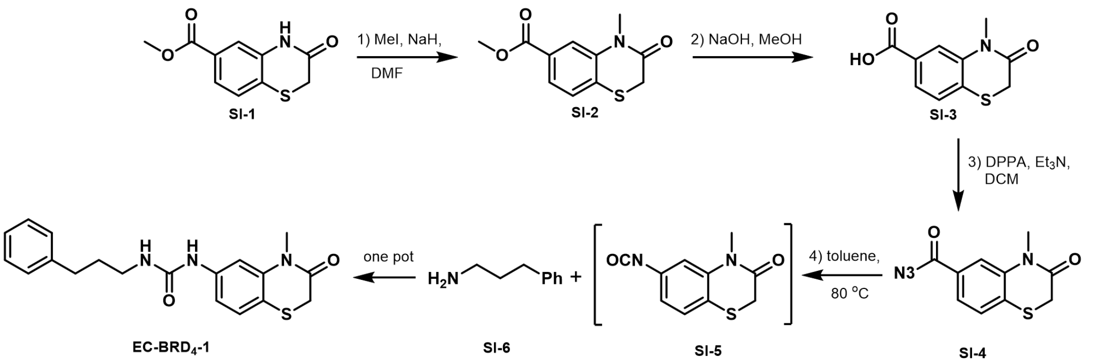


To a stirring solution of **SI-1** in DMF, NaH was added portion-wise at 0 °C under argon. The reaction mixture was then stirred for an additional 10 minutes at this temperature before the addition of MeI. After 20 minutes, the reaction was quenched by the slow addition of water at 0°C. Afterward, the mixture was extracted by ethyl acetate three times and the combined organic phases were washed with Brine once and dried over $Na_2SO_4$. After the concentration of the organic phase under reduced pressure, the resulting residue was purified by column chromatography on silica gel (PE/EA = 6/1) to give pure product **SI-2** as a yellow solid.

The **SI-2** was dissolved in MeOH followed by the addition of NaOH in $H_2O$ at room temperature. The resulting mixture was stirred for 10 hours before it was acidified to PH = 3 with 3 M aqueous HCl. The reaction was extracted with ethyl acetate three times, and the combined organic phases were washed with Brine once, dried over $Na_2SO_4$, filtered, and concentrated under reduced pressure. The crude white solid product acid **SI-3** was directly used in the next step without further purification.

$Et_3N$ and diphenylphosphoryl azide (DPPA) were added to a stirring solution of acid **SI-3** in DCM at room temperature. After 4 hours, the solvent was directly dried on the rotary evaporator. The resulting residue was purified by column chromatography packed with silica gel with PE/EA = 8/1 as the eluent to give the desired acyl azide **SI-4**

The **SI-4** was heated to 80 °C in toluene and stirred for 4 hours to form intermediate isocyanate **SI-5**. Then the heating oil bath was removed to cool down the reaction to room temperature, which was followed by the addition of amine **SI-6**. The mixture was then stirred overnight at room temperature. (white suspension formed) After the removal of the toluene in a vacuum, the white solid residue was purified by the silica gel-packed column chromatography, while eluting with PE/EA = 1/1 to the desired product **EC-BRD4-1** as a white solid.

$^{1}$H NMR (400 MHz, $CDCl_3$) δ 7.42 (d, *J* = 2.1 Hz, 1H), 7.31 – 7.27 (m, 1H), 7.23 – 7.13 (m, 4H), 6.72 (dd, *J* = 8.3, 2.1 Hz, 1H), 6.59 (s, 1H), 3.40 (s, 3H), 3.36 (s, 2H), 3.29 (t, *J* = 7.1 Hz, 2H), 2.67 (t, *J* = 7.6 Hz, 2H), 1.87 (p, *J* = 7.3 Hz, 2H), δ 1.65 (s, 2H).

$^{13}$C NMR (101 MHz, $CDCl_3$) δ 166.1, 155.6, 141.4, 141.1, 138.3, 128.7, 128.6, 128.5, 126.2, 117.4, 115.0, 110.1, 40.2, 33.4, 32.3, 31.8, 31.7.

**Synthesis of EC-BRD4-2**

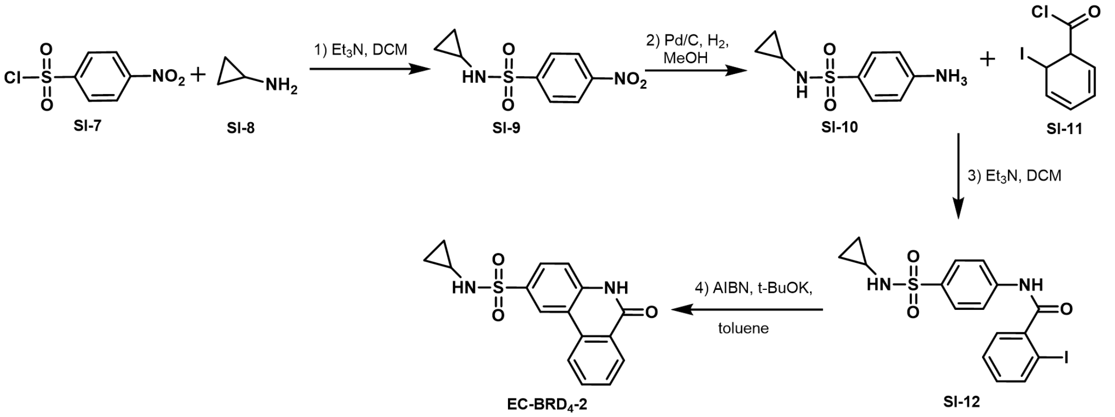


The amine **SI-8** (243 μL, 3.50 mmol, 1.00 eq.) and $Et_3N$ (2.44 mL, 17.5 mmol, 5.00 eq.) were dissolved in $CH_2Cl_2$ (10.0 mL) at room temperature under argon. Then the sulfonyl chloride **SI-7** (854 mg, 3.85 mmol, 1.10 eq.) was added and the reaction was stirred for 4 hours before it was directly dried on the rotary evaporator. The crude product was eluted with PE/EA = 2/1 on a silica gel-packed column to give product **SI-9** as a white solid in a quantitive yield.

The nitro-compound **SI-9** (100 mg, 0.413 mmol) was dissolved in MeOH (4.10 mL) at room temperature under argon. The Pd/C (10% Pd on charcoal, 20.0 mg) was added in one portion under argon. Afterward, the flask was pumped with a water pump and refilled with $H_2$ ballon for three cycles. The reaction was stirred overnight and filtered through a short pad of Celite. After concentration under reduced pressure, the corresponding aniline **SI-10** was used for the next reaction without further purification.

To a solution of crude aniline compound **SI-10** (161 mg, 0.836 mmol, 1.00 eq.) in $CH_2Cl_2$ (8.4 mL) was added acyl chloride **SI-11** (245 mg, 0.920 mmol, 1.10 eq.) at room temperature under argon. The reaction was stirred overnight and quenched with water. The mixture was then extracted with $CH_2Cl_2$ three times. The organic phase was dried over $Na_2SO_4$, filtered, and concentrated under reduced pressure. The resulting residue was purified by a silica gel-packed column and eluted with PE/EA = 2/1 to 1/1 to give the desired product **SI-12** (70.8 mg, 19%) as a white solid.

The **SI-12** (20.0 mg, 0.045 mmol, 1.00 eq.), AIBN (1.50 mg, 0.009 mmol, 0.200 eq.), and *t*-BuOK (25.0 mg, 00.225 mmol, 5.00 eq.) were added to a dry Schlenk flask under argon, followed by the addition of toluene (2.00 mL) at room temperature. Then the reaction was heated to 85 °C and stirred for 30 hours. Then, the reaction was extracted with EA three times. The organic phase was dried over $Na_2SO_4$, filtered, and concentrated in vacuo. Further purification with column chromatography on silica gel (elution: PE/EA = 2/1 to 1/1) afforded the desired product **EC-BRD4-2**.

**Synthesis of EC-V2R-1**

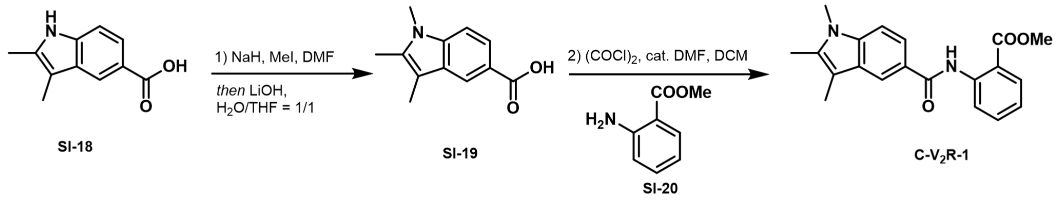


The commercially available compound **SI-18** (150 mg, 0.800 mmol, 1.00 eq.) was dissolved in anhydrous DMF (4.00 mL) at 0 °C, followed by the addition of NaH (60 % dispersion in mineral oil, 96.0 mg, 2.40 mmol, 3.00 eq.). After stirring for 10 minutes at room temperature, the MeI (454 mg, 3.20 mmol, 4 eq.) was added to the above clear reaction mixture, and the reaction mixture was stirred for an additional 30 minutes at room temperature to methylate the free indole and the carboxylic acid. To hydrolyze the methyl ester, THF (4.00 mL), $H_2O$ (4.00 mL), and LiOH (76.8 mg, 3.20 mmol, 4.00 eq.) were added successively to the above mixture at room temperature. After stirring overnight, the reaction mixture was diluted with EA and $H_2O$. Then the reaction mixture was washed with aqueous NaOH solution (1.00 M) twice, the basic aqueous phases were collected and acidified by adding aqueous HCl solution (3.00 M) to PH = 4. The resulting white suspension was extracted with EA twice. The combined organic phase was washed with Brine once, dried over $Na_2SO_4$, filtered, and concentrated under reduced pressure. The crude product **SI-19** was directly used in the next step without purification.

The acid **SI-19** (110 mg, 0.541 mmol, 1.00 eq.) was loaded in a dry Schlenk flask under argon, followed by the addition of DMF (2.00 μL, 0.027 mmol, 0.050 eq.) and DCM (5.40 mL) to give a white suspension. After cooling to 0 °C, the oxalyl chloride (60.0 μL, 0.704 mmol, 1.30 eq.) was added. The reaction was warmed to room temperature by the removal of the ice bath and stirred till the reaction mixture was clear. Then the aniline compound **SI-20** (180 μL, 1.41 mmol, 2.6 eq.) was added and the resulting reaction mixture was stirred overnight to give a grey suspension. The reaction mixture was diluted with water and extracted with a large amount of EA. The combined organic phase was dried over $Na_2SO_4$, filtered, and concentrated in vacuo. The flash column chromatography on silica gel (elution: PE/EA = 4/1) gave the desired product **EC-V2R-1** as a white solid.

$^1$H NMR (400 MHz, $CDCl_3$) δ 12.03 (s, 1H), 9.00 (dd, *J* = 8.5, 1.2 Hz, 1H), 8.28 (d, *J* = 1.8 Hz, 1H), 8.08 (dd, *J* = 8.0, 1.7 Hz, 1H), 7.85 (dd, *J* = 8.6, 1.8 Hz, 1H), 7.61 (ddd, *J* = 8.7, 7.2, 1.7 Hz, 1H), 7.31 (d, *J* = 8.6 Hz, 1H), 7.12 – 7.06 (m, 1H), 3.97 (s, 3H), 3.68 (s, 3H), 2.37 (s, 3H), 2.32 (s, 3H).

$^{13}$C NMR (101 MHz, $CDCl_3$) δ 169.1, 167.4, 142.6, 138.7, 134.8, 134.4, 131.0, 128.5, 125.4, 122.1, 120.6, 119.8, 118.9, 115.2, 108.5, 108.1, 52.5, 29.9, 10.4, 8.9.

**Synthesis of EC-V2R-2**

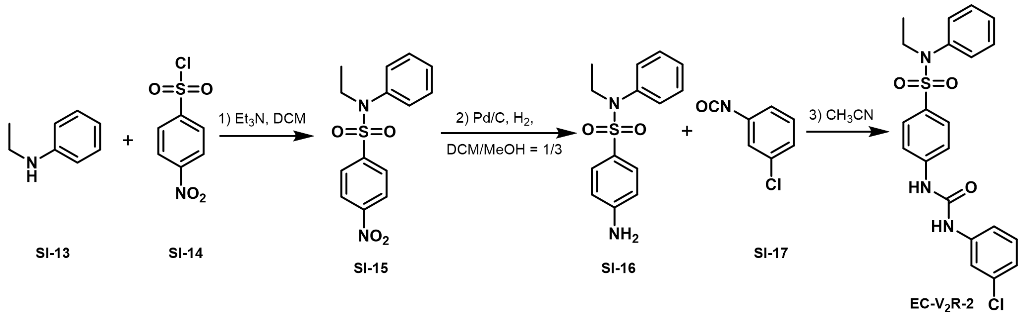


The aniline **SI-13** (1 mL, 7.94 mmol, 1.00eq.) and $Et_3N$ (1.66 mL, 11.9 mmol, 1.50 eq.) were mixed in the round-bottom flash at room temperature under argon. The sulfonyl chloride **SI-14** (2.11 g, 9.53 mmol, 1.20 eq.) was added portion-wise at the same temperature. When the TLC indicates the absence of **SI-13**, the reaction was quenched by saturated aqueous $NaHCO_3$ solution and extracted with $CH_2Cl_2$ three times. Afterward, the combined organic phase was dried over $Na_2SO_4$, filtered, and concentrated under reduced pressure. The crude product **SI-15**

was directly used in the next step without further purification.

All obtained crude **SI-15** was redissolved in $CH_2Cl_2$/MeOH (1/3, 7.00 mL/21.0 mL) at room temperature under argon. After the addition of Pd/C (10% Pd on charcoal, 200 mg), the inner argon was changed to $H_2$ gas via three pumping-refilling cycles. The reaction was stirred overnight and filtered through a short pad of Celite. After concentration under reduced pressure, the corresponding aniline **SI-16** was used for the next reaction without further purification.

The commercially available 3-chlorphenylisocyanate **SI-17** (87.0 μL, 0.720 mmol, 1.20 eq.) was mixed with **SI-16** (167 mg, 0.600 mmol, 1.00 eq.) in $CH_3CN$ (3 mL) at room temperature. The reaction was stirred overnight before the reaction solvent was removed under reduced pressure. The residue was purified by column chromatography on silica gel, eluting with PE/EA = 1/1 to the desired product **EC-V2R-2** as a white solid.

$^1$H NMR (400 MHz, $CDCl_3$) δ 7.50 – 7.43 (m, 3H), 7.38 (d, *J* = 8.5 Hz, 2H), 7.34 – 7.28 (m, 3H), 7.25 – 7.18 (m, 2H), 7.13 – 6.96 (m, 3H), 3.58 (q, *J* = 7.1 Hz, 2H), 1.07 (t, *J* = 7.1 Hz, 3H); $^{13}$C NMR (101 MHz, $CDCl_3$) δ 152.4, 143.5, 139.4, 138.4, 134.8, 131.0, 130.2, 129.4, 129.1, 129.0, 128.6, 123.8, 119.9, 118.6, 117.9, 46.0, 14.1.

### S6 Bioassay Methods

**Cell Culture.** Followed by previous studies[9], HEK293 cells expressing the SNAP-tagged human vasopressin V2R receptor cells were maintained in Dulbecco's modified Eagle's medium (Thermo Fisher, Shanghai) containing 10% (v/v) newborn calf serum (Gibco, Shanghai), streptomycin (100 μg/mL), penicillin (100 IU/mL), and G418 (0.6 mg/mL) at 37 °C in 5% CO2. The cells were subcultured twice a week.

**Equilibriumand Kinetic Binding Experiments.** Compound EC-V2R-1 and EC-V2R-2 were tested in the presence of a 6.3 nM fluorescent ligand at 37°C. The affinity was tested in the presence of unlabeled benzodiazepine derivatives at various concentrations. By adding SNAP-tagged V2R cells (6000 cells/well) to each well, the experiment was initiated. Signals were recorded after 1h of incubation at 37°C. Nonspecific conivaptan-red binding was measured with 0.3 μM unlabeled conivaptan. The fluorescence intensity and the HTRF ratios were calculated as described previously[9].

**Fluorescence Resonance Energy Transfer Assay.** Compound EC-BRD4-1 and EC-BRD4-2 were subjected to HTRF binding assay to detect their binding ability to the BRD4-BD1 domain. The BRD4-BD1 inhibitor screening assay kit (EPIgeneous™ Binding Domain kit A, no. 62BDAPEG) was purchased from Cisbio Bioassays. Biotin-labeled histone H4 (no. AS-64989-1) was purchased from AnaSpec Inc., and BRD4 protein (no. RD-11-157) was purchased from Cisbio Bioassays. The HTRF signal in this assay is correlated with the amount of biotin-labeled histone binding to bromodomain, and the final concentration of DMSO is 1% in all reactions. All binding reactions were conducted at room temperature. The 20 μL reaction mixture in the assay buffer contains BRD4-BD1 (4 μL), biotin-labeled histone (4 μL), compound (2 μL), streptavidin-d2 conjugate (5 μL), and anti-GST-Eu3+ cryptate conjugate (5 μL). For the negative control (blank), 6 μL of the assay buffer was added instead of the protein and compound, and for the positive control, 2 μL of the assay buffer was added instead of the compound. The reaction mixture was incubated for 3 hours. After the incubation, the HTRF signal was measured using the Synergy™H1 configurable multi-mode microplate reader.

1 Hinze, S. R. *et al.* Beyond ball-and-stick: Students' processing of novel STEM visualizations. *Learning and instruction* **26**, 12-21 (2013).
2 Sebens, C. T. Electron charge density: a clue from quantum chemistry for quantum foundations. *Foundations of Physics* **51**, 75 (2021).
3 Skogh, M. *et al.* The electron density: a fidelity witness for quantum computation. *Chemical Science* **15**, 2257-2265 (2024).
4 Nakatsuji, H. Electron-cloud following and preceding and the shapes of molecules. *Journal of the American Chemical Society* **96**, 30-37 (1974).
5 Leckband, D. & Israelachvili, J. Intermolecular forces in biology. *Quarterly reviews of biophysics* **34**, 105-267 (2001).
6 Kaufman, B. *et al.* COATI: Multimodal contrastive pretraining for representing and traversing chemical space. *Journal of Chemical Information and Modeling* **64**, 1145-1157 (2024).
7 Barducci, A., Bonomi, M. & Parrinello, M. Metadynamics. *Wiley Interdisciplinary Reviews: Computational Molecular Science* **1**, 826-843 (2011).
8 Grimme, S. Exploration of chemical compound, conformer, and reaction space with meta-dynamics simulations based on tight-binding quantum chemical calculations. *Journal of chemical theory and computation* **15**, 2847-2862 (2019).
9 Liu, C. *et al.* Revisit ligand-receptor interaction at the human vasopressin V2 receptor: A kinetic perspective. *European Journal of Pharmacology* **880**, 173157 (2020).
